## Supplementary Information for "amr.watch – monitoring antimicrobial resistance trends from global genomics data"

### Supplementary Material

#### Supplementary Methods

*Retrieval of WGS data and associated metadata from INSDC databases*

We retrieve all metadata associated with pathogens in the 2017 WHO priority pathogen list [1] from the European Nucleotide Archive (ENA) via the ENA Portal API on an ongoing basis. We proceed with entries (samples) from the relevant pathogens (based on the submitted classification; see **Supplementary Table 1** for accepted taxon IDs) with an available collection date from 2010 onwards that is decodable to at least the year, as well as a sampling location decodable to at least the country level. For the latter, we attempt to assign an ISO 3166-1 alpha-2 country code [2] based on information available in either the "location", "lat"/"long" or "country" metadata fields (assessed in this order). We use the 249 "officially assigned" country codes and "other" code types where necessary (currently "XK" (Kosovo) and "AN" (Netherlands Antilles)). Reverse geocoding of latitude and longitude coordinates is performed using two open-source Node.js packages: geojson-places [3] and geojson-geometries-lookup [4] with a high quality GeoJSON map of countries' political maritime borders with 5m resolution [5]. Geocoding of location names from the "country" field is performed using the OpenCage Geocoding API [6] and the Mapbox Geocoding API [7], and samples are discarded if the resulting country codes from the two tools are inconsistent. A manually-curated list of non-standard location names and their associated country codes is also maintained and applied where necessary. Additionally, we use an additional Node.js package, country-to-iso [8], to handle inconsistent country names. Finally, collection dates (found in many different formats among the ENA metadata) are converted to a standard ISO 8601 format where they can be unambiguously interpreted. Entries that have collection dates with ranges that cover more than one year are disregarded. Only the year is extracted when the available format renders the precise date ambiguous (i.e. when the month/day values are both ≤12 and the month and day values are not equal, such as 01/02/2013).

Of those entries with a decodeable country code and collection date, we proceed with those with library_strategy="WGS", library_source="GENOMIC", library_layout="PAIRED" and instrument_platform= "ILLUMINA". If an entry is associated with more than one sequencing run, the run with the highest number of total bases is selected using the "base_count" field. We exclude sequencing runs that are linked to more than one entry (sample). Sequencing runs with only one fastq file are also disregarded. Sequencing runs must possess at least 20X mean coverage, based on assessing the total number of bases ("base_count") relative to the expected length of the genome (**Supplementary Table 1**).

Sequence reads fulfilling the above criteria are downloaded from the Sequence Read Archive (SRA) using SRA-Toolkit fastq-dump v3.1.0 [9] after checking that they exist in the SRA using Entrez Programming Utilities [10].

*Genome assembly*

Sequence reads are assembled with a workflow [11] that uses the SPAdes assembler v3.15.3 [12] and which is implemented in Nextflow v21.04.1.5556 (versions as of March 2025) [13]. The workflow includes steps for quality control (QC) of the reads pre- and post-trimming, and determination of key QC metrics of the final assemblies using QUAST v5.0.2 (version as of March 2025) [14].

*Species verification*

The Speciator tool (v4.0.0) within Pathogenwatch [15] is used to verify the species of the genome assemblies. The SISTR tool (v1.1.1) [16, 17], implemented within Pathogenwatch [18], is additionally used to assign the serotype of *Salmonella enterica* genomes. Genome assemblies with a species and/or serotype identification that do not match defined taxon IDs for each pathogen (**Supplementary Table 1**) are excluded. Note that for each of *E. coli*, *S. sonnei* and *S. flexneri*, we accept genomes annotated as either *E. coli* or *Shigella* in the ENA, with the Speciator assignments used in subsequent processing (thus permitting inconsistencies in this case).

*QC of genome assemblies*

Genome assemblies that are identical to another assembly in the ongoing curated collection are excluded on the basis that these represent duplicate uploads of the same raw sequence data to INSDC databases. Assemblies are also excluded if they fail to meet one or more of our defined QC criteria, which are specific to each species/serovar (**Supplementary Table 1**). Defined thresholds for the number of contigs and N50 values were developed based on manual inspection of the species-specific distributions from all genomes assembled up to March 2022. The range of accepted GC content is the observed range among all RefSeq genomes of each species/serovar. The range of accepted assembly length values were determined using the known range among RefSeq genomes, allowing for an additional ±5%.

*Variant typing*

We use Pathogenwatch to identify the sequence type (ST) of all pathogens via available multilocus sequence typing (MLST) schemes from PubMLST [19], except for *Salmonella enterica subsp. enterica* serovar Typhi and *Streptococcus pneumoniae* (see below). For assemblies belonging to *Acinetobacter baumannii*, the "Pasteur" MLST scheme rather than "Oxford" scheme is used due to the presence of *gdhB* paralogues in the latter [20].

Pathogens utilising other typing schemes with amr.watch are *Salmonella enterica subsp. enterica* serovar Typhi, for which we use GenoTyphi [21] as implemented in Pathogenwatch, and *Streptococcus pneumoniae* for which we use Global Pneumococcal Sequencing Cluster (GPSC) assignments [22] as also implemented in Pathogenwatch.

*Identification of markers associated with antimicrobial resistance*

We identify genes and mutations associated with antimicrobial resistance (AMR) in each quality-controlled assembly using AMRFinderPlus v3.10.23 and database v2021-12-21 (versions as of March 2025) [23]. AMR markers are identified for antimicrobial classes defined in the 2017 WHO priority pathogen list [1], with the addition of those for quinolone resistance in *E. coli*. The list of AMR determinants included for each pathogen-antimicrobial combination is available in **Supplementary Table 2** and has been developed based on a comprehensive literature review. In particular, we disregard some markers reported by AMRFinderPlus for which there is insufficient experimental validation of their phenotypic outcome. Only complete matches to AMR markers are accepted.

#### Supplementary Figures

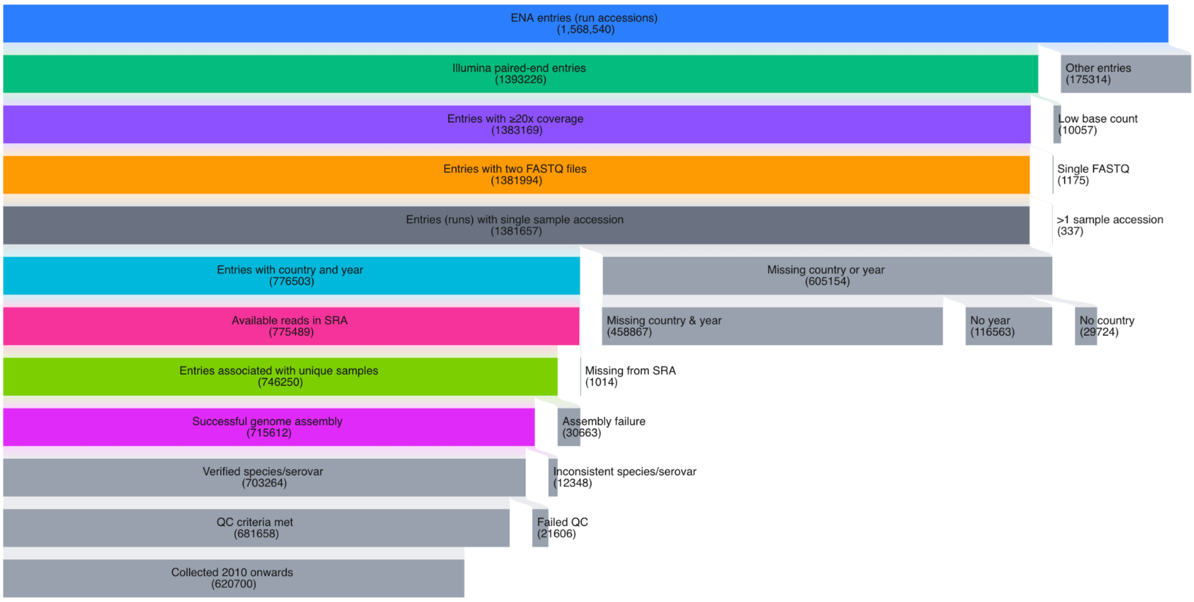


**Supplementary Figure 1.** Overview of the filtering process within the amr.watch workflow applied to all public genomes of priority bacterial pathogens available in the International Nucleotide Sequence Database Collaboration (INSDC) databases up to 31 March 2025. The same visualisation, updated in real-time, can be viewed at: https://amr.watch/summary/all Individual visualisations for each pathogen can also be accessed from: https://amr.watch/summary
